# Estimating global variation in the maximum growth rates of eukaryotic microbes from cultures and metagenomes via codon usage patterns

**DOI:** 10.1101/2021.10.15.464604

**Authors:** JL Weissman, Edward-Robert O. Dimbo, Arianna I. Krinos, Christopher Neely, Yuniba Yagües, Delaney Nolin, Shengwei Hou, Sarah Laperriere, David A. Caron, Benjamin Tully, Harriet Alexander, Jed A. Fuhrman

## Abstract

Microbial eukaryotes are ubiquitous in the environment and play important roles in key ecosystem processes, including accounting for a significant portion of global primary production. Yet, our tools for assessing the functional capabilities of eukaryotic microbes in the environment are quite limited because many microbes have yet to be grown in culture. Maximum growth rate is a fundamental parameter of microbial lifestyle that reveals important information about an organism’s functional role in a community. We developed and validated a genomic estimator of maximum growth rate for eukaryotic microbes, enabling the assessment of growth potential for organisms and communities directly in the environment. We produced a database of over 700 maximum growth rate predictions from genomes, transcriptomes, and metagenome-assembled genomes. By comparing the maximal growth rates of existing culture collections with environmentally-derived genomes we found that, unlike for prokaryotes, culture collections of microbial eukaryotes are only minimally biased in terms of growth potential. We then extended our tool to make community-wide estimates of growth potential from over 500 marine metagenomes, mapping growth potential across the global oceans. We found that prokaryotic and eukaryotic communities have highly correlated growth potentials near the ocean surface, but there is no correlation in their genomic potentials deeper in the water column. This suggests that fast growing eukaryotes and prokaryotes thrive under similar conditions at the ocean surface, but that there is a decoupling of these communities as resources become scarce deeper in the water column.

## Introduction

Microbial eukaryotes are ubiquitous in the environment, and play diverse roles relevant to ecosystem (e.g., [1, 2]) and human (e.g., [3, 4]) health. In the oceans in particular, protists dominate, accounting for approximately 30% of total marine biomass and of primary producer biomass as well [5]. Marine systems account for about half of all global primary production [6], hence the abundance of protists in these systems indicates a central role for protists in regulating global biogeochemical cycles [7]. And yet, our tools for studying the ecology and evolution of eukaryotic microbes are still quite limited, at least in comparison to their prokaryotic relatives [8].

Several recent developments have greatly advanced our ability to survey the ecology of microbial eukaryotes directly from the environment using metagenomics. Large-scale efforts to augment the sizes of our existing genomic and transcriptomic databases, specifically the Marine Microbial Eukaryote Transcriptome Sequencing Project (MMETSP; [9]), have expanded our ability to use database-dependent approaches for metagenomic analysis for both taxonomic and functional classification (e.g., [10–13]). At the same time, novel approaches for binning and validation have been applied by multiple groups to reconstruct high-quality metagenome-assembled genomes (MAGs) from environmental datasets [14–16].

With these new environmentally-derived genomes come new challenges – specifically that of inferring features of an organism’s physiology and ecology from its genome sequence, a persistent challenge in metagenomics [17–19]. One trait of particular interest is the maximal growth rate of an organism, a fundamental parameter of microbial lifestyle that can tell us a great deal about an organism’s ecology [20–22]. Among microbial eukaryotes, minimal doubling times range over two orders of magnitude, from hours (e.g., [23, 24]) to days (e.g., [25]), and potentially even weeks (e.g., [26, 27]). Temperature sets a well-studied upper-bound on the maximal growth rates of microbial eukaryotes (see work on the Eppley Curve, e.g., [28–33]), but there is a great deal of variation among species below this threshold.

For prokaryotes, genomic signals of translation optimization can be leveraged in order to predict the maximal growth rates of an organism [20, 22, 34]. Here, we show that the same signals, specifically the codon usage bias of highly expressed genes, can be used to estimate the growth potential of eukaryotic microbes, and of entire mixed-species communities, directly from their genome or metagenome sequences. We compiled a database of 178 species of eukaryotic microbes with recorded growth rates in culture and either publicly available genomes or transcriptomes. We then used this database to build a genomic predictor of growth potential for eukaryotic microbes. We applied this tool to a set of 465 MAGs and 517 metagenomes to derive ecological insights about the global variation of eukaryotic growth potential across organisms and environments.

## Results and Discussion

### Predicting maximal growth rates of eukaryotic microbes

We compiled a database of maximal growth rates and optimal growth temperatures recorded in culture for 178 species with either genomic or transcriptomic information publicly available (S1 Fig, S1 Table). A sizeable portion of this dataset corresponded to marine eukaryotic microbes, with 101 entries corresponding to organisms in the MMETSP, though eukaryotic microbes from other environments were also represented, including important human pathogens (e.g., *Giardia intestinalis*, *Entamoeba histolytica*, *Leshmania spp.*, etc.). In general, eukaryotic microbes with genomes in GenBank, which tended to be human-associated, had faster maximal growth rates than the marine eukaryotic microbes found in MMETSP (S1 and S2 Fig). This pattern is similar to that found in prokaryotes, where human-associated bacteria and archaea typically had much faster growth rates than those found in marine systems [20].

One of the most reliable signals of optimization for rapid growth in prokaryotic genomes is high codon usage bias (CUB) in highly expressed genes [22]. The degeneracy of the genetic code means that multiple codons may code for the same amino acid, but not all organisms use alternative codons at equal frequencies. In fact, many organisms, both prokaryotic and eukaryotic, are biased in their usage of alternative codons. There are many forces acting on codon bias – including mutational biases and selection for mRNA stability [35] – but the codon usage patterns of genes are also thought to be optimized to the relative abundance of tRNAs within the cell to facilitate rapid translation. This optimization is particularly apparent among highly-expressed genes in fast-growing species [36–43]. The basic intuition is that the optimization of genes for rapid translation leads to increased CUB, and in turn facilitates rapid growth. We wanted to see whether such patterns can be leveraged to predict the growth rates of eukaryotic microbes [44]. For each genome or transcriptome, we calculated the CUB of a set of highly expressed genes relative to the expected CUB calculated from all other coding sequence in that genome or transcriptome (details of these calculations can be found in Materials and Methods; [20, 40, 45]). Because ribosomal proteins are expected to have high expression across species and many physiological conditions [20, 22], we take these as our set of highly expressed genes for all analyses (see Methods and S2 Table). We found a significant negative relationship between CUB of highly expressed genes calculated in this way and the minimum doubling time of an organism (Pearson correlation with Box-Cox transformed doubling times, *ρ* = −0.47, *p* = 5.64 × 10^−11^). Thus, we confirmed that high CUB is a signal of growth rate optimization among microbial eukaryotes.

We then built a linear model relating CUB of highly expressed genes and optimal growth temperature to the doubling times of eukaryotic microbes (Fig 1a; S2 Fig). Optimal growth temperature has been seen to be an important parameter predicting the maximum growth rates of prokaryotes that is important to control for in genomic predictors [20, 22]. We found that our model could explain about a third of variation in doubling time among organisms (linear regression, *r*^2^ = 0.337), and that both CUB (*p* = 4.48 × 10^−8^) and optimal growth temperature (*p* = 3.18 × 10^−8^) were significant predictors in the model. Interestingly, similar to what we have previously reported for prokaryotes [20], we saw that for eukaroytic microbes, the relationship between CUB and doubling time may saturate at a threshold doubling time, after which CUB no longer changes with increasing doubling time (Fig 1b). In our earlier work on prokaryotes, we took the presence of this threshold as evidence that for slow-growing organisms who experienced little selection for optimized codon usage, drift would overwhelm the evolutionary process [20]. It appears that a similar dynamic may be at work among eukaryotes, although the small number of sequenced eukaryotic microbes with long minimal doubling times (greater than 30-40 hours) makes this hard to assess. If such a threshold does exist for eukaryotes, it is at a much higher doubling time than the one seen in prokaryotes (30-40 hours versus 5 hours respectively; [20]), likely due to constraints on eukaryotic growth related to cell size and complexity. It is possible that predicted maximum growth rates for very slow growing organisms will be overestimated using our tool, though we lack strong evidence that this will be the case. To determine pragmatic cutoffs for users, we redefined the doubling time cutoffs for reliable prokaryotic and eukaryotic prediction in terms of the CUB rather than the predicted doubling times to get temperature-independent cutoffs appropriate for organisms growing in colder environments (see Methods). A clear limitation of our approach is that we can never say for sure whether an organism’s experimentally measured maximum growth rate is its actual maximum growth rate, which may in part explain why organisms with slow empirical growth measurements sometimes have faster maximum growth rates inferred genomically.

**Fig 1.**
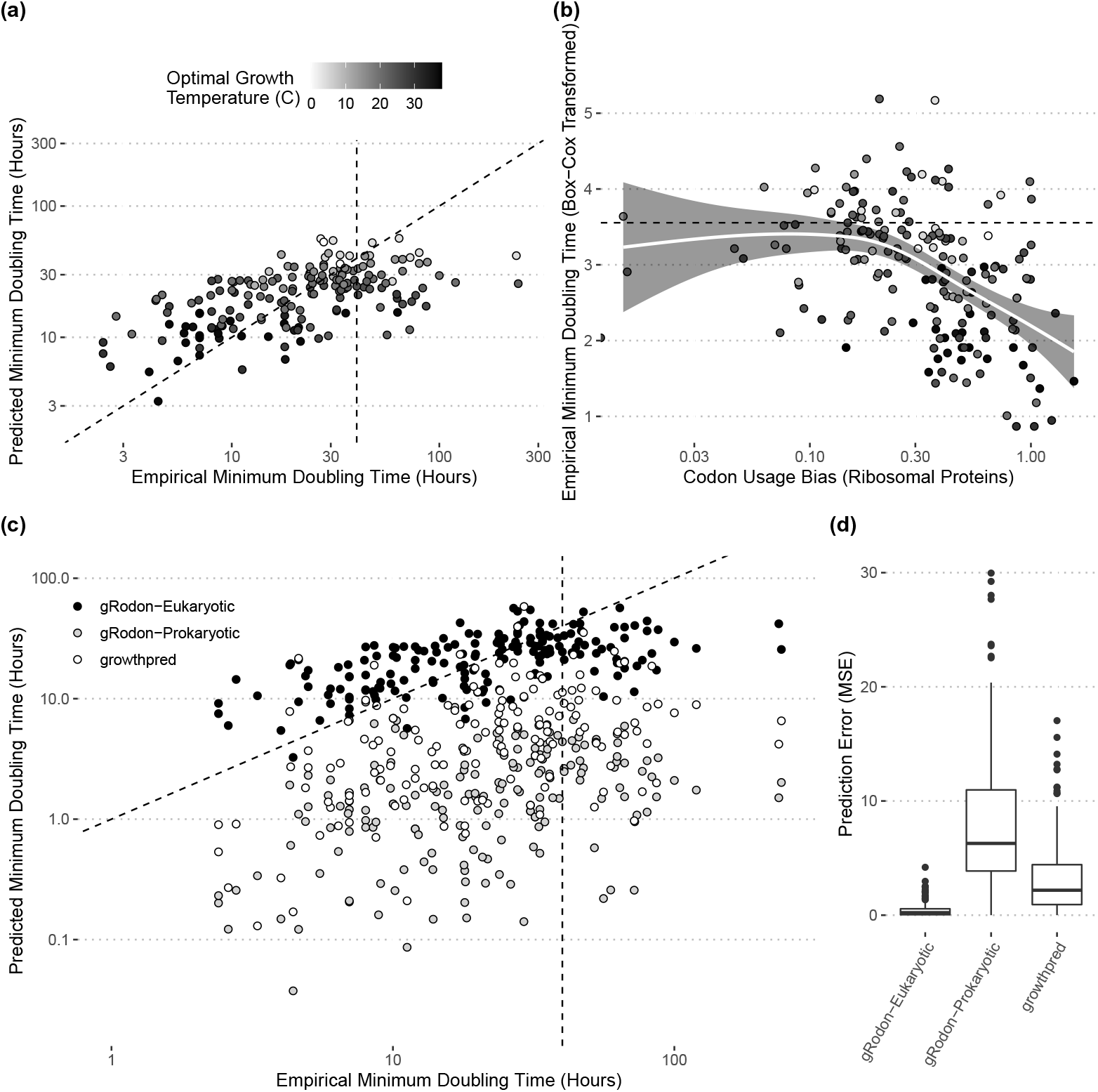
Codon usage bias (CUB) and temperature predict the maximum growth rates of microbial eukaryotes. (a) Predictions from a linear model of minimum doubling time with CUB and temperature as predictors on our training set generally reflect empirically observed doubling times (*r*^2^ = 0.328). (b) The relationship between CUB and minimum doubling time is roughly linear and negative until approximately 30-40 hours (dashed line at 40 hours; Pearson correlation, *ρ* = −0.48, *p* = 3.74 × 10^−9^), after which the relationship levels off (Pearson correlation, *ρ* = −0.099, *p* = 0.55). Gray line is a smoothing line generated with ggplot2. (c) Predictions of the maximum growth rates of microbial eukaryotes on the basis of CUB and temperature using models trained on prokaryotes are systematically biased towards faster growth predictions and (d) perform much worse than a model trained directly on eukaryotes in terms of mean squared error (MSE). Dashed vertical lines denotes 40 hours and dashed diagonal line denotes where *x* = *y*.

We asked whether our eukaryotic model of growth improved predictions for eukaryotic organisms relative to the predictions made by previous tools developed for and trained exclusively on prokaryotes [20, 22]. We applied these prediction tools to our eukaryotic dataset and found that they systematically overestimated the growth rates of eukaryotic organisms, often by more than an order of magnitude, indicating poor performance on eukaryotes (Fig 1c-d). Thus, our eukaryote-specific model is an important development, as prokaryote-specific models cannot accurately predict eukaryotic growth rates. We have incorporated this eukaryote-specific model into the open-source and user-friendly growth prediction R package gRodon, which we previously developed to predict prokaryotic growth rates (https://github.com/jlw-ecoevo/gRodon; [20]).

Finally, we considered the possibility that our model was overfit to the data, which would mean our predictor would perform poorly on new datasets. Overfitting is a particularly relevant concern when dealing with species data where shared evolutionary history can induce a hierarchical dependency structure between datapoints. In addition to traditional cross-validation, we applied a blocked cross-validation approach, which controls for phylogenetic structure by taking each phylum in our dataset as a fold to hold out for independent error estimation rather than holding out random subsets of our data as in traditional cross-validation. Even under these controls, eukaryotic gRodon showed a clear improvement over prokaryotic models as well as against prediction using the average growth rate of a group (S3 and S4 Figs). This suggests CUB-growth relationships extrapolate well across phyla. Given that the genomes taken from NCBI and transcriptomes from MMETSP are from very different sets of microbes, we wondered whether the choice of data source would impact our model. We trained models on each source dataset and saw good cross-prediction between training datasets (S5 Fig).

### Environmentally derived genomes reveal biases in culture collections and ecological patterns

We obtained a large set of 1669 eukaryotic MAGs assembled and binned from the Tara Oceans metagenomes by two separate groups [15, 16]. Of these, we were able to predict the growth rates of 465 MAGs in which we found at least 10 ribosomal proteins (see Materials and Methods for details). These MAGs were uniformly slow-growing, with an average minimum doubling time of approximately one day, and none with a minimum doubling time less than 10 hours long (Fig 2a). These MAGs provide a baseline expectation of the maximal growth rates of eukaryotic microbes in marine environments, and while the reconstruction of MAGs from the environment is not a wholly unbiased process, we expect these MAGs to be more representative of the distribution of organisms living in the environment than what we find within our culture collections [20]. In fact, we found that MAGs were estimated to have similar, though marginally distinguishable, doubling times to cultured organisms in MMETSP (means of 26.1 and 25.1 hours respectively; Kolmogorov-Smirnov test, *p* = 0.0187; Fig 2a). The differences between these two datasets were most apparent when looking at the tails of the distributions of growth rates, where the MMETSP had a long tail of fast-growing organisms that was absent among the MAGs (Fig 2a,b). Altogether the data suggest that our culture databases of eukaryotes do a good job at capturing the distribution of growth rates among organisms, though they are slightly enriched for fast growing organisms that are relatively rare in the environment. This result is in stark contrast to the pattern seen among marine prokaryotic organisms, where culture collections were shown to be systematically biased towards fast-growers [20]. A caveat to these analyses is that while MAGs are likely to better approximate the natural distribution of organisms in the environment than culture collections, the binning process used to construct MAGs may also be biased, meaning that MAGs do not perfectly represent natural populations of microbes. That being said, we found only extremely weak (and likely spurious) associations between completeness and inferred growth rate across MAGs (and no or an opposite pattern was seen for relative abundance to that of growth rate; S6 Fig), suggesting no strong association between the ability to bin an organism’s genome and its maximum growth rate.

**Fig 2.**
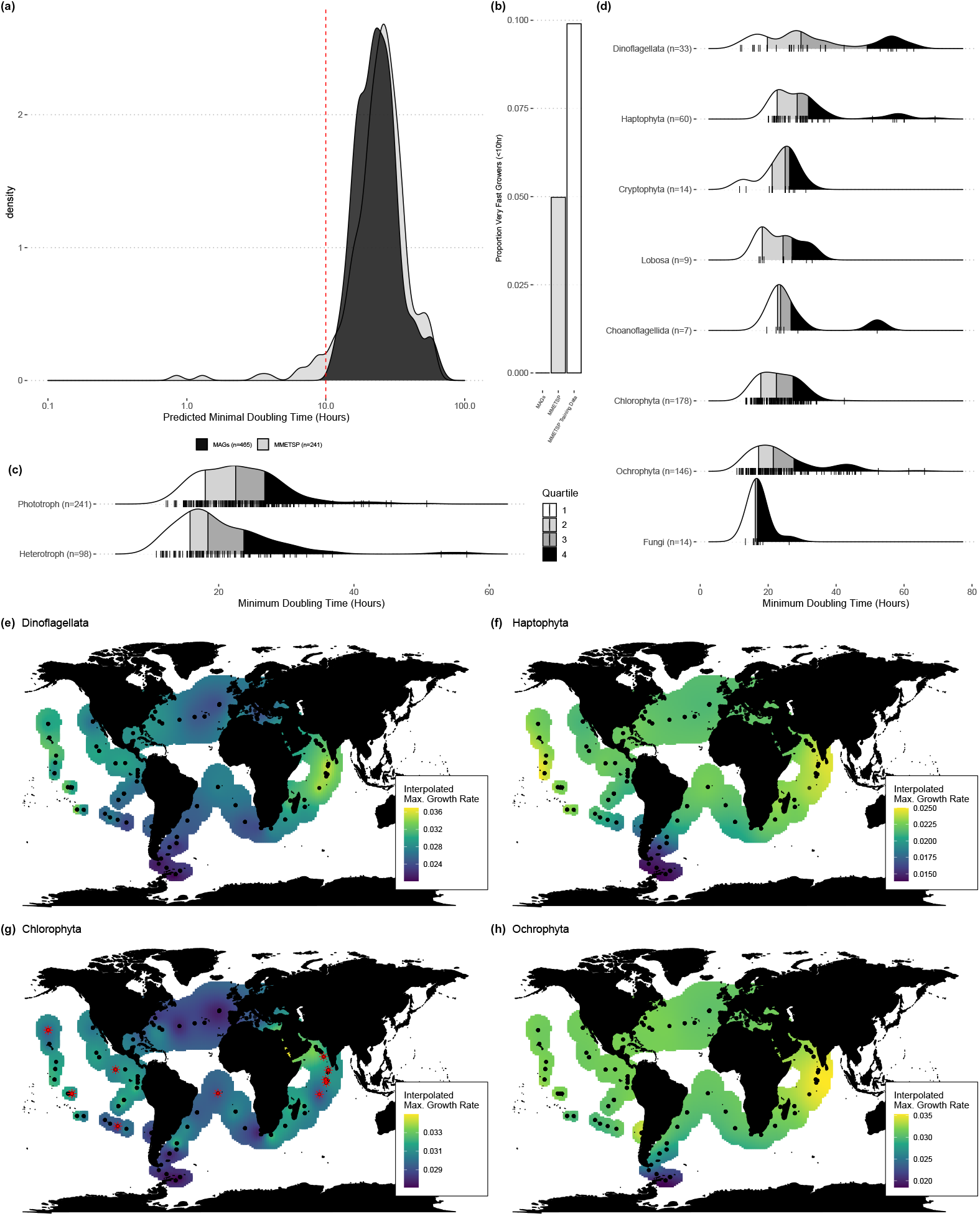
Environmentally derived genomes of eukaryotic microbes reveal differences in growth potential across sampling sources, lifestyles, and taxonomic groups. (a) The distribution of predicted minimum doubling times of organisms represented in the MMETSP is nearly indistinguishable from the distribution of predicted minimum doubling times among marine eukaryotic microbes represented by MAGs. (b) Even so, there are small number of very fast growing organisms (minimum doubling time < 10 hours) in MMETSP that form a long tail absent in the MAG data. (c) MAGs from organisms predicted to be heterotrophic were associated with faster maximum growth rates than those predicted to be phototrophic. (d) Different taxonomic groups have distinct distributions of predicted growth potentials among their members, as predicted from MAGs. (e-h) Using the abundance of MAGs from groups with many representatives mapped back to the Tara Oceans surface metagenomes (< 100 meters) we found group-specific patterns in average maximum growth rate (h^−1^) across the world’s oceans. Red asterisks denote possible overestimates due to saturation of the CUB-growth relationship. For panels (a-d) see S7 Fig for an analysis of possible overestimated rates, which cannot account for observed patterns.

Within the set of MAGs, several patterns were apparent. First, while organisms classified as phototrophic and heterotrophic had largely overlapping growth rate distributions (Fig 2c), heterotrophs tended to grow faster than phototrophs (Mann-Whitney U Test, *p* = 7.65 × 10^−6^; trophic classification on the basis of the presence of metabolic pathways in a MAG, taken from Alexander et al. [15]). This reflects previous findings that at higher temperatures heterotrophic marine eukaryotic microbes had faster growth rates than phototrophic ones, though phototrophs outgrew heterotrophs at lower temperatures because their growth rates decreased less dramatically with decreases in temperature [33].

Just as growth rates varied among functional groups, they also systematically varied among taxonomic groups (Fig 2d). Overall, marine fungi had the fastest average estimated growth rates. MAGs belonging to the Chlorophyta also seemed to be relatively fast growing, with a somewhat narrow range of growth rates clustered around a doubling time of about a day. By contrast Dinoflagellata, Haptophyta, and to some degree Ochrophyta all had a considerable number of very slow growing representatives (minimal doubling time > 40 hours), though these groups had very broad distributions of growth rates and included many faster growing members as well. The diversity of growth rates in these groups is perhaps not surprising, as the cell sizes of diatom and dinoflagellate species vary over two orders of magnitude, indicating a wide diversity of morphologies and environmental niches [46, 47]. Overall, the distribution of maximal growth rates varied across taxonomic groups, likely a product of both specialization for different niches and historical contingency.

Finally, we looked at the maximum growth rate and abundance of each MAG across the Tara Oceans dataset [15, 48] to get group-specific maximum growth rates across the global oceans. We found the average maximum growth rates of several major groups of microbial eukaryotes with many representative MAGs by taking an abundance-weighted maximum growth rate for each Tara surface sample (Fig 2e-h; < 100 meters). Groups varied in their global growth patterns, but in general the Indian Ocean was a hotspot for organisms with high maximum growth rates and the Southern Ocean was dominated by organisms with low maximum growth rates. Ochrophyta (Fig 2f) and Dinoflagellata (Fig 2e) communities varied widely in their growth rates, whereas Chlorophyta (Fig 2g) community maximum growth rates varied less across samples. Haptophyta (Fig 2h) communities generally had the lowest maximum growth rates seen across samples.

### Predicting the growth potential of prokaryotic and eukaryotic communities from metagenomes

It is often difficult to reconstruct high-quality MAGs for many organisms, both prokaryotic and eukaryotic, from the environment. Even when we cannot easily obtain MAG representatives of every community member in a particular environment, it is possible to apply CUB-based predictors to a metagenome to estimate the average maximum growth rate of that community [22, 49]. The prokaryotic growth predictor previously implemented in the gRodon package allowed the user to predict the median community-wide maximal growth rate of the prokaryotic community [20], and we recently updated and benchmarked metagenome mode v2 for improved community-level prediction [49]. Our eukaryotic model can be similarly applied to calculate the median maximal growth rate of the eukaryotic community represented in a metagenomic sample (see Methods for details). We benchmarked this approach against synthetic mixtures of genomes meant to simulate mixed-species communities and saw good performance predicting community-wide averages, in line with our previous results for prokaryotes (see S8-S9 Figs and Methods).

To demonstrate this application, we obtained assemblies of 610 globally-distributed marine metagenomic samples from the BioGEOTRACES dataset [50]. This dataset is particularly useful for our purposes because samples were not size-fractionated, allowing both prokaryotic and eukaryotic communities to be assessed simultaneously. Overall, we were able to predict the average community-wide maximal growth rates of the prokaryotic and eukaryotic communities in 517 samples with at least 10 ribosomal proteins each that could be classified as eukaryotic or prokaryotic (Fig 3; S10 Fig). The correlation between the growth potentials of prokaryotic and eukaryotic communities was striking (Pearson correlation of samples from < 100 meters, *ρ* = 0.457, *p* = 1.74 × 10^−27^; Fig 3a-b), though this relationship varied with depth (Fig 3a-d). A linear model confirmed a significant interaction between depth and the relationship between eukaryotic and prokaryotic growth rates (linear regression of prokaryotic growth rates, *β*_eukaryotes_ = 0.185, *p*_eukaryotes_ = 2.81 × 10^−11^, *β*_depth_ = 4.78 × 10^−4^, *p*_depth_ = 2.02 × 10^−3^, *β*_eukaryotes:depth_ = 1.53 × 10^−4^, *p*_eukaryotes:depth_ = 0.0120). Notably, these correlations were not simply byproducts of temperature shifts, as the CUB of eukaryotic and prokaryotic communities co-varied across surface samples (though correlations among deeper samples could be explained by temperature; Fig 3c-d). Additionally, these patterns could noy be attributed to differences in coverage. While doubling time did decrease with the relative abundance of eukaryotic contigs in a sample, as would be expected, samples with a lower proportion of eukaryotes were not particularly skewed in their estimated growth rates (S11 Fig).

**Fig 3.**
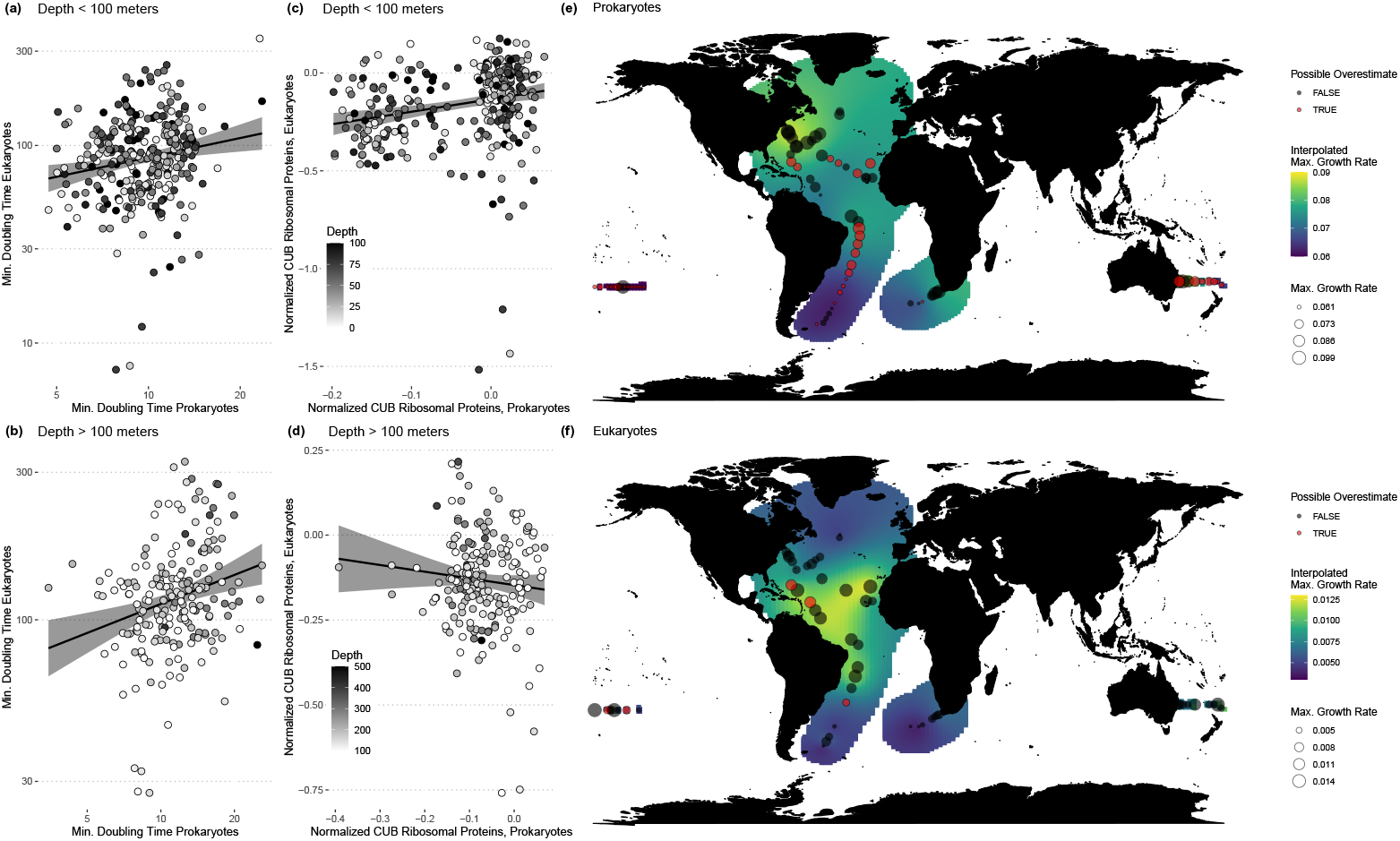
Bulk community-wide prediction of growth potential from metagenomes yields insights into global patterns of eukaryotic and prokaryotic growth in the oceans. (a-b) The average community-wide maximum growth rate of eukaryotic and prokaryotic communities are correlated near the ocean surface (< 100 meters; *ρ* = 0.240, *p* = 2.48 × 10^−5^), and at deeper depths (> 100 meters; *ρ* = 0.457, *p* = 5.83 × 10^−12^). (c-d) Yet this correlation is at least partly driven by temperature, particularly for deeper samples. While (c) the normalized codon usage bias of eukaryotic and prokaryotic communities are correlated at the surface (< 100 meters; *ρ* = 0.223, *p* = 7.70 × 10^−5^), (d) they are not at depth (> 100 meters; *ρ* = −0.0884, *p* = 0.208). (e-f) Average community-wide growth rates near the surface for eukaryotes and prokaryotes vary substantially across the global oceans (< 100 meters) and, while associated, have distinct overall patterns.

It is perhaps not surprising that the growth potentials of eukaryotic and prokaryotic communities would be correlated, since conditions favorable to more copiotrophic lifestyles (e.g., high nutrients) should be similar across both prokaryotes and eukaryotes. The observed decoupling of the CUB of eukaryotic and prokaryotic communities with increasing depth aligns with a shift towards increasingly heterotophic eukaryotic communities (S13). Prokaryotic communities at the surface have correlated CUB with deeper prokaryotic communities, but eukaryotic communities are not correlated across depths, nor with prokaryotic communities at other depths (S12 Fig). Altogether, this suggests that any physical processes linking the surface to deeper depths (e.g., sinking particles) may occur on a different timescale to processes shaping the community growth potential (e.g., community assembly).

## Conclusions

We developed and validated a new tool to estimate the growth potential of eukaryotic microbes directly from genomic, transcriptomic, and metagenomic sequences. Using this tool, we were able to predict the maximal growth rates of a large set of uncultured marine organisms directly from reconstructed MAGs. We found distinct patterns in growth potential across functional and taxonomic groups and assessed existing culture collections for functional bias. We then applied our tool to a large set of marine metagenomes to predict the community-wide growth potential of eukaryotes along large ocean transects. We found a clear positive relationship between eukaryotic and prokaryotic growth potential at the ocean surface, suggesting that fast growing organisms from multiple domains of life thrive under similar conditions, and the same for slow growing organisms. With an increasing number of environmental metagenomes published each year, for many environments it will now be possible to build high-resolution maps of microbial growth potential across domains, yielding insights into the drivers of microbial community structure and function.

We emphasize that gRodon estimates the maximum growth rate of an organism or the average maximum growth rate of a community, not the instantaneous growth rate at any given time. In the wild, it is likely that microbes rarely reach these maximum rates. Nevertheless, this parameter gives us an idea about the ecological role of an organism, in particular whether it has undergone selection for the ability to replicate rapidly. Microbial ecologists often conceptualize organisms’ growth strategies on a spectrum from a “boom-bust” strategy favoring rapid maximum growth rates versus a “slow-and-steady” strategy with slower maximum growth rates, and call organisms with these two strategy sets “copiotrophs” and “oligotrophs” respectively [20, 21, 51]. In our previous work on prokaryotes, we found that these distinct growth classes have different evolutionary patterns, gene content, and functional profiles [20]. When applied at the community level, the average community-wide maximum growth rate can be thought of as an “index of copiotrophy” which measures the relative frequency of these two distinct growth classes in the community [49].

Our tool demonstrates the clear utility of genomic and metagenomic trait estimators for eukaryotic microbes. Yet, when working with eukaryotic microbes there are relatively few bioinformatic resources both in terms of methods and databases. Moving forward, as the complexity and subtlety of our bioinformatic tool-set increases, eukaryotic microbes represent a new frontier for methods development and ecological investigations with molecular data (e.g., [11, 12, 14–16]).

## Materials and Methods

The code to generate all figure and analysis in the paper can be found at https://github.com/jlw-ecoevo/eeggo. The new gRodon v2 R package with the eukaryotic growth rate model implemented can be found at https://github.com/jlw-ecoevo/gRodon2. All visualizations were made using the ggplot2 [52] and ggpubr [53] R packages. For mapping, we used the maps package [54], as well as the automap package for spatial interpolation (universal blocked kriging with the autoKrige() function; [55]). Ridgeline plots were generated using R package ggridges [56].

See S1.1 Text for supplemental methods describing how we generated the datset used to train our model (and the datasets and programs used [11–13, 33, 57–64]).

See S1.2-1.3 Text and S14-S15 Figs for supplemental methods describing the processes for predicting maximum growth rates from MAGs and Metagenomes using eukaryotic gRodon (and the datasets and programs used [10–12, 14–16, 20, 49, 50, 65–71]).

### Fitting the Model

For each MMETSP transcriptome in our training dataset we used the annotations provided [58] (generated using dammit [72]) to locate coding sequence corresponding to ribosomal proteins. For each GenBank genome in our training dataset we searched among translated coding sequences for ribosomal proteins using blastp v2.10.1 [65] against a custom blast database of ribosomal proteins of eukaryotic microbes drawn from the Ribosomal Protein Gene Database (all genes coding for ribosomal proteins available from Dictyostellium discoideum, Giardia lamblia, Phaeodactylum tricornutum, *Plasmodium falciparum, Thalassiosira pseudonana*, and *Toxoplasma gondii*; S2 Table; [66]). In all downstream analyses we omitted any genomes or transcriptomes with fewer than 10 ribosomal proteins detected [20, 22].

For each coding sequence corresponding to a ribosomal protein in each genome or transcriptome we calculated the MILC (Measure Independent of Length and Composition) statistic of CUB [40] using the coRdon R package [45], the same as done for prokaryotic gRodon [20]. This statistic is both GC-content and length corrected and should be insensitive to both factors. For these calculations the expected codon usage was taken as the genome-wide average (across all coding sequences in a genome or transcriptome; [20]). As recommended in the coRdon documentation, in order to get a reliable estimate of codon bias we removed all genes with fewer than 80 codons. We then calculated the median CUB across all genes coding for ribosomal proteins for each genome or transcriptome.

We then fit a linear model to Box-Cox transformed doubling times (with the optimal *λ* chosen using the boxcox() function from the MASS package [73]) using (1) optimal growth temperature, and (2) the normalized median CUB of genes coding for ribosomal proteins

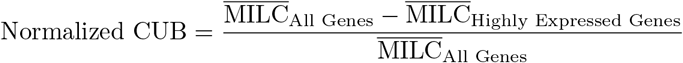

(see [22, 49] for details of normalization procedure) as predictors. We then implemented this model into the existing gRodon package for prokaryotic growth rate prediction, expanding the package’s predictive range to eukaryotic organisms (using the new mode=“`eukaryotes`” setting; https://github.com/jlw-ecoevo/gRodon2). For gRodon’s eukaryotic metagenome mode the codon-usage bias was calculated separately on a per-gene basis prior to normalization, as done with prokaryotic metagenome mode v2 [49].

For comparison with prokaryotic models we ran genomes and transcriptomes through growthpred (obtained as a docker image at https://hub.docker.com/r/shengwei/growthpred; [74]) and gRodon v1.0.0 on metagenome mode (the prokaryotic setting most similar to both growthpred and our eukaryotic model; [20, 22]), including the recorded optimal temperatures for prediction.

Appropriate CUB cutoffs for prediction were determined by taking a maximum growth rate model trained only on mesophilic organisms (optimal growth temperature between 20C and 60C), but otherwise not accounting for temperature (otherwise as above). We then determined at what CUB value the model predicted a doubling time of 30 hours (approximately where the CUB vs growth relationship saturates; Fig 1b; normalized CUB of 0.012 for eukaryotes, a similar procedure with a 5 hour threshold identified a CUB cutoff of 0.59 for the original prokaryotic model [20]). These cutoffs can be used to determine whether a maximum growth estimate from gRodon is likely to be an overestimate, independent of the growth temperature of an organism (e.g., Fig 3e-f).

## Supporting information

Supplemental References

## Acknowledgments

J.L.W. was supported by a postdoctoral fellowship in marine microbial ecology from the Simons Foundation (Award 653212). A.I.K. was supported by the Computational Science Graduate Fellowship (DOE; DE-SC0020347). H.A was supported by a National Science Foundation grant (OCE-1948025). We also acknowledge support from Simons Foundation Collaboration on Computational Biogeochemical Modeling of Marine Ecosystems (CBIOMES) Grant 549943 (to J.A.F.) and US NSF Division of Ocean Sciences (OCE) Grant 1737409 (to J.A.F.).

