## Supplemental References for "Estimating global variation in the maximum growth rates of eukaryotic microbes from cultures and metagenomes via codon usage patterns"

[68] Josef Konrad. *In vitro culture of* Babesia divergens*:: biochemical and immunological investigations*. PhD thesis, Imperial College London, 1986.

[82] Sala Manuel. *Effect of temperature on growth and production of YTXs and lytic compounds by* Protoceratium reticulatum *from Greenland*. PhD thesis, EMBC, Uni Bremen, 2014.

[137] Eliăka Trsková. Mechanism of photoprotection in photosynthetic proteins. Master's thesis, University of South Bohemia in České Budějovice, Faculty of Science, České Budějovice, 2015.
