## Supplemental Figures and S2 Table for "Estimating global variation in the maximum growth rates of eukaryotic microbes from cultures and metagenomes via codon usage patterns"

### Supplementary Figures and Tables - Estimating the maximal growth rates of eukaryotic microbes from cultures and metagenomes via codon usage patterns

JL Weissman, Edward-Robert O. Dimbo, Arianna Krinos, Christopher Neely, Yuniba Yagües, Delaney Nolin, Shengwei Hou, Sarah Laperriere, David A. Caron, Benjamin Tully, Harriet Alexander, Jed A. Fuhrman

October 24, 2022

#### SText 1 Supplemental Methods

##### SText 1.1 Training Data

From a list of protist species with transcriptomes in MMETSP and/or genomes in GenBank, we searched for recorded growth rates in culture, alongside the temperatures at which these maximum rates were recorded, from the scientific literature (S1 Fig, S1 Table). If multiple rates were reported in culture, we always took the fastest rate we were able to find. We converted between doubling time and specific growth rate using the equation: doubling time $= \frac{\ln(2)}{\text{growth rate}}$. When growth rate was reported in terms of divisions per day we instead used the conversion equation: doubling time $= \frac{1}{\text{growth rate}}$. A considerable number of growth rates recorded for marine organisms came from previous database compilation efforts by others [1, 2].

All MMETSP transcriptome assemblies with annotated coding sequence were obtained from `https://zenodo.org/record/3247846` [3]. Species that had a eukaryotic prey species listed under experimental conditions were excluded from further analysis to reduce the possibility of contamination. For each species in MMETSP, we selected the single largest transcriptome assembly. We then removed any potential cross-contamination between species by identifying possible contaminants using CroCo v1.1 [4] with a threshold of 97% identity and otherwise default parameters (`--suspect-id 97`), removing any transcripts listed as “suspect”. We additionally classified transcripts using kraken2 v2.1.1 [5, 6] with the ‘nt’ database and default parameters, and removed any transcripts classified as viruses or prokaryotes in order to remove any potential contaminants.

For assemblies from GenBank with growth rates listed in our dataset we first ran EukMetaSanity v1.0.0 [7], which incorporates repeat prediction [8], reference protein selection [9, 10], and ab-initio gene predictions to determine putative gene loci. The output GeneMark-EP/ES predictions were used for analysis [11,12]. We then classified coding sequences using kraken2 v2.1.1 [5,6] with the ‘nt’ database and default parameters, and removed any transcripts classified as viruses or prokaryotes in order to remove any potential contaminants.

#### SText 1.2 Estimating Growth Rates from MAGs

We obtained a set of 1669 eukaryotic MAGs assembled from the Tara Oceans metagenomic surveys by two groups [13, 14]. Previously, EukMetaSanity had been run on these MAGs to call genes and find coding sequences [7]. We used these annotations, and (similar to above) searched for ribosomal proteins using blastp v2.10.1 [15] against a custom blast database of ribosomal proteins of eukaryotic microbes drawn from the Ribosomal Protein Gene Database [16]. We ran EUKulele v1.0.6 to classify these MAGs taxonomically and omitted any organisms identified as Metazoa from downstream analyses [17]. Division-level classifications were taken as the division assigned as most likely by eukulele. After removing any MAGs with less than 10 ribosomal proteins detected or that were classified as Metazoa, we were left with a total of 465 eukaryotic MAGs.

To infer the optimal growth temperatures of each MAG we used distributional data across the Tara Oceans metagenomes. For MAGs from Delmont et al. [13], optimal temperatures had already been predicted by the authors as part of a machine-learning pipeline implemented to discover each MAGs niche. For the Alexander et al. [14] MAGs we took a simpler approach. For each MAG we took the top 1% of samples in terms of MAG relative abundance and calculated the mean temperature recorded for those samples (S14 Figure). For closely related MAGs found in both Delmont et al. [13] and Alexander et al. [14], we found that the two methods for estimating optimal growth temperature agreed well (S15 Figure).

Finally, we calculated maximal growth rate using gRodon v2.0.0 in eukaryote mode.

For the Tara Ocean analysis, TPM values and taxonomic classifications for each MAG in each sample were obtained from Alexander et al. [14] for all MAGs in that dataset. Average maximum growth rates for each sample-taxonomy combination were calculated as the weighted median of the MAGs from that taxonomic group present in that sample, with TPM used as the MAG weights.

#### SText 1.3 Estimating Growth Rates from Metagenomes

Assemblies of the bioGEOTRACES metagenomes were obtained from Biller et al [18]. We then ran EukRep v0.6.6 on these assemblies to classify contigs as eukaryotic or prokaryotic (using settings `-m strict --tie prok`; [19]).

In order to call and annotate genes from prokaryotic contigs we ran prokka v1.14.6 (using settings `--norrna --notrna --metagenome --cpus 80 --centre X --compliant`; [20]). Ribosomal proteins were identified from prokka annotations. We then predicted the average community-wide maximal growth rate of the prokaryotic community using gRodon v2.0.0 in metagenome mode using the temperature metadata provided with the samples [21].

In order to call genes from eukaryotic contigs we ran MetaEuk v4.a0f584d (using setting `easy-predict`; [9]). Similar to our procedure with individual genomes, we searched for ribosomal proteins among translated proteins output from MetaEuk using blastp v2.10.1 [15] against a custom blast database of ribosomal proteins of eukaryotic microbes drawn from the Ribosomal Protein Gene Database [16]. We then predicted the average community-wide maximal growth rate of the eukaryotic community using gRodon v2.0.0 in eukaryote mode using the temperature metadata provided with the samples.

In order to benchmark eukayotic gRodon's ability to predict the average maximal growth rates of mixed-species eukaryote communities we generated several community-level datasets (see [22] for more detail and justification of this benchmarking procedure for mixed-species communities). To start, we generated three sets of 10,000 "gene mixtures" drawn from two sources: (1) genomes of eukaryotic microbes from GenBank, and (2) MMETSP transcriptomes. We distinguish "gene mixtures" from true synthetic metagenomes, where mixtures are simply the concatenated set of coding sequences from the constituent genomes or transcriptomes. In contrast, a "true" synthetic metagenome would be produced via the generation of synthetic reads from genomic references. By limiting ourselves to gene mixtures, we focus our analysis on issues of prediction on mixed communities only, and do not consider possible complications arising from the sequencing or assembly process (though see [22] for community-level benchmarking with true synthetic metagenomes). To simulate a gene mixture we randomly sampled pairs genomes from the relevant set of genomes and concatenated annotated coding sequences from these genomes into a single fasta file for that mixture. This was repeated 10,000 times for each data source. Maximum growth rates were inferred for all source genomes using gRodon's "full" eukaryotic prediction mode to derive ground-truth against which to benchmark predictions on the genome mixtures. In this way, community wide benchmarking tests whether community-level predictions recapitulate individual level predictions, but do not directly address accuracy of these predictions on natural communities (for which independent maximum growth rate measurement is not currently possible).

##### SText 1.4 Phylogeny

We ran BUSCO v5.2.2 against the `eukaryota_odb10` database on translated proteins from the complete set of MAGs described above as well as the decontaminated MMETSP transcriptomes [23]. For the purposes of tree-building only, we removed any organisms without at least 50% of BUSCO gene families present (out of 255). We then identified any gene families present in at least 80% of remaining organisms, yielding 51 gene families. We aligned each of these families using MUSCLE v3.8.31 with default parameters [24] and trimmed our alignments with trimal v1.4.rev15 (using setting `-automated1`; [25]). Alignments were concatenated (using `https://github.com/nylander/catfasta2phyml`) and trimmed again using trimal v1.4.rev15 (using setting `-automated1`). We then used Fasttree v2.1.10 to infer a phylogeny (using default settings; [26]).

We visualized our phylogeny using R package ggtree [27] and calculated patristic distance using R package ape [28].

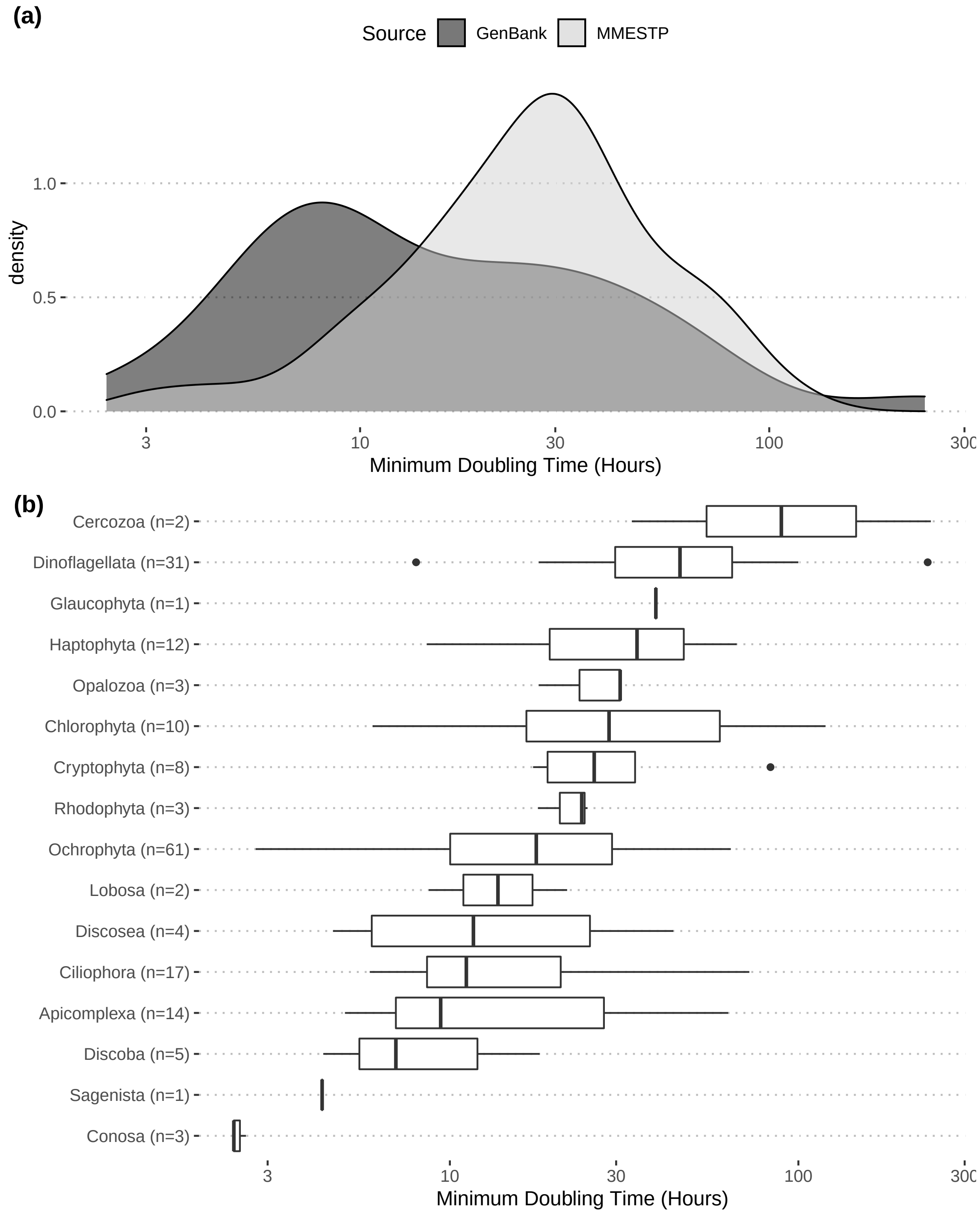


Figure S1: Distributions of recorded minimum doubling times for 178 organisms taken from the literature and used to train our model ($n = 101$ for organisms with transcriptomes in MMETSP and $n = 77$ for organisms with genomes in GenBank). (a) Note that organisms in GenBank (including many pathogens) in general had faster growth rates than the marine organisms in MMETSP (t-test on log-transformed values, $p = 0.0001854$). (b) The taxonomic breakdown of the training set.

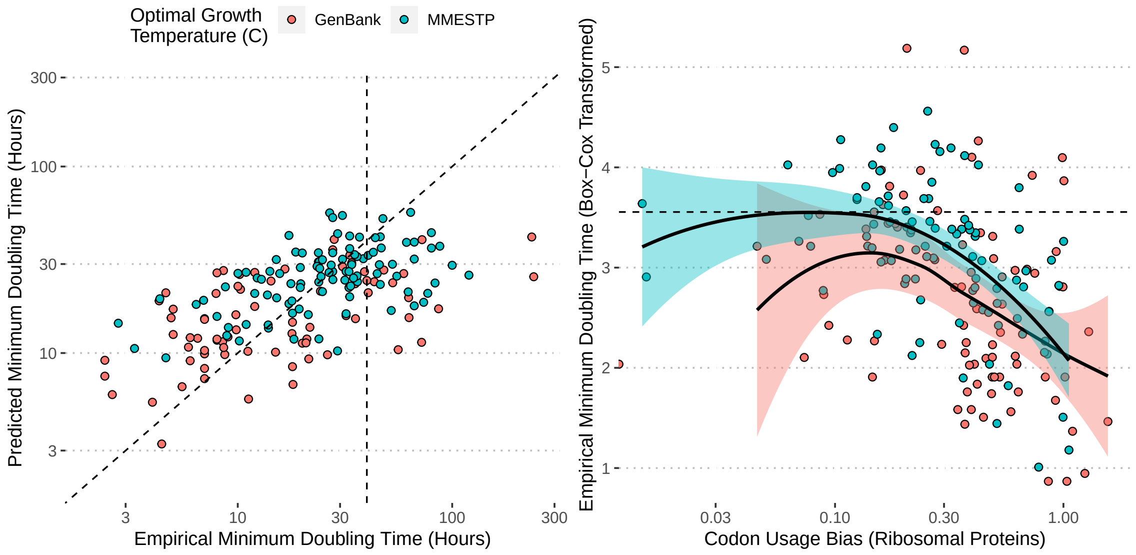


Figure S2: Figure 1 panels a-b broken down by source database.

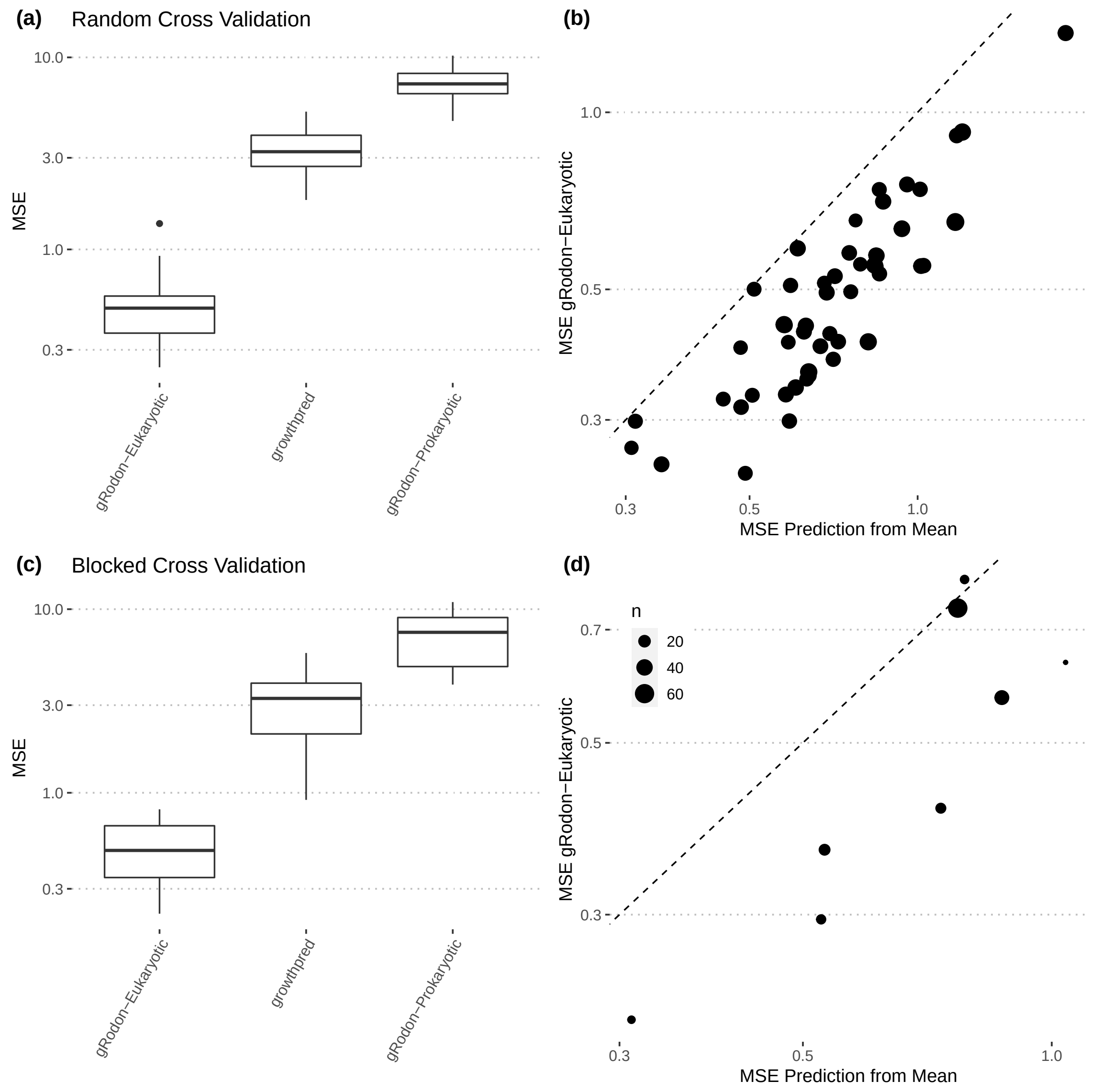


Figure S3: Cross validation confirms eukaryotic gRodon outperforms other models on eukaryotic organisms. (a) Cross validation confirms the results of Fig 1c-d. (b) Comparison of gRodon predictions against predictions generated by taking the average growth rate of the training folds in box-cox transformed space and extrapolating this to held out folds. (c) Blocked cross validation using phyla as folds confirms the results of Fig 1c-d. (b) Comparison of gRodon predictions against predictions generated by taking the average growth rate of the training set in box-cox tranformed space and extrapolating this to held out folds (where folds are phyla).

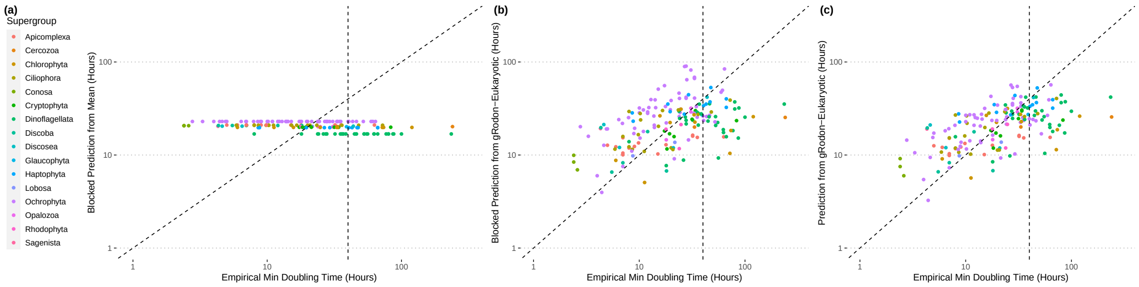


Figure S4: Actual versus predicted doubling times for each phylum in the training data set using (a) the average growth rate of all other phyla in box-cox transformed space, (b) the eukaryotic gRodon model trained on all other phyla (holding that phylum out), and (c) the full eukaryotic gRodon model.

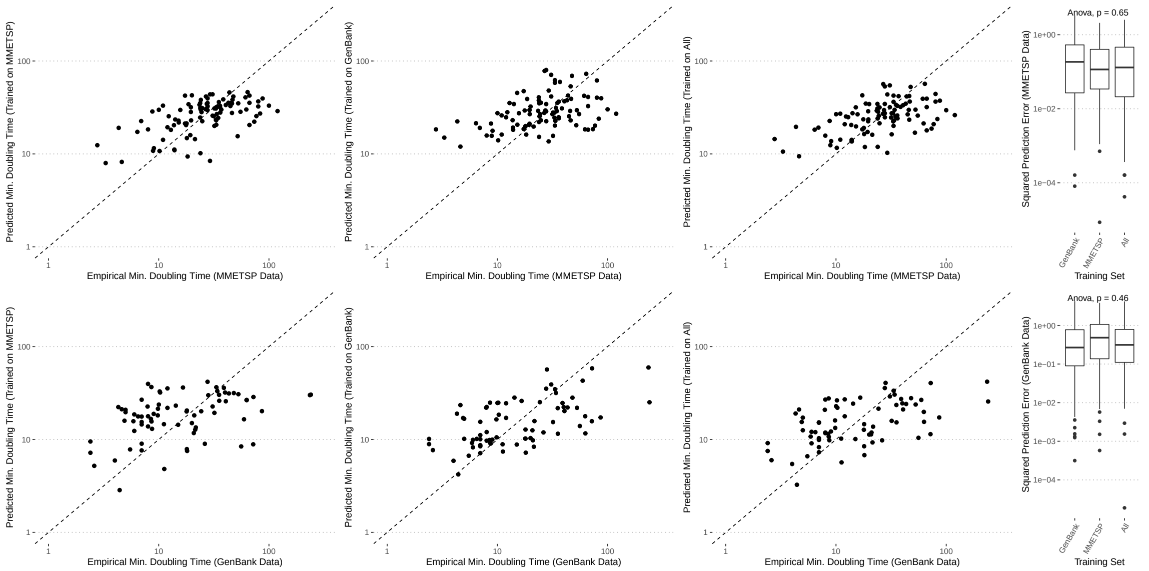


Figure S5: The gRodon model works well across training sets. The model trained on MMETSP data predicts well across test data sets, and the model trained on data from GenBank performs similarly well across test data.

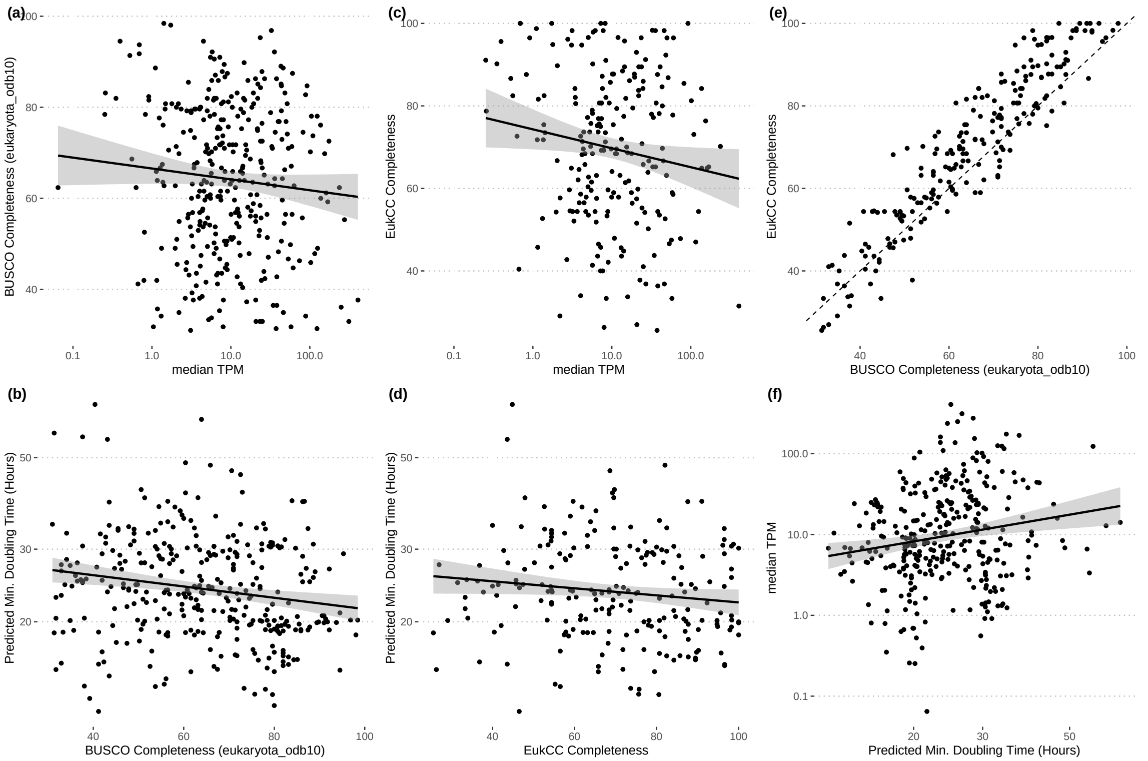


Figure S6: MAG completeness varies weakly with abundance and inferred maximum growth rate across Tara Oceans samples. (a-b) BUSCO completeness did not vary significantly with MAG abundance (linear regression, $p = 0.109$, $r^2 = 0.00465$), but did with predicted minimum doubling time (linear regression, $p = 6.11 \times 10^{-4}$, $r^2 = 0.0312$), though very weakly. (c-d) EukCC completeness did vary significantly with MAG abundance (linear regression, $p = 0.0317$, $r^2 = 0.0167$), as well as with predicted minimum doubling time (linear regression, $p = 0.038$, $r^2 = 0.0153$), albeit very weakly. (a-d) These relationships are quite possibly spurious and have essentially no explanatory power.(c) The two completeness metrics used agree well (Pearson correlation, $\rho = 0.935$, $p < 10^{-16}$). (d) Surprisingly, abundance was negatively associated with max growth rate across MAGs, though this relationship was again extremely weak and possibly spurious (linear regression, $p = 1.43 \times 10^{-4}$, $r^2 = 0.0390$).

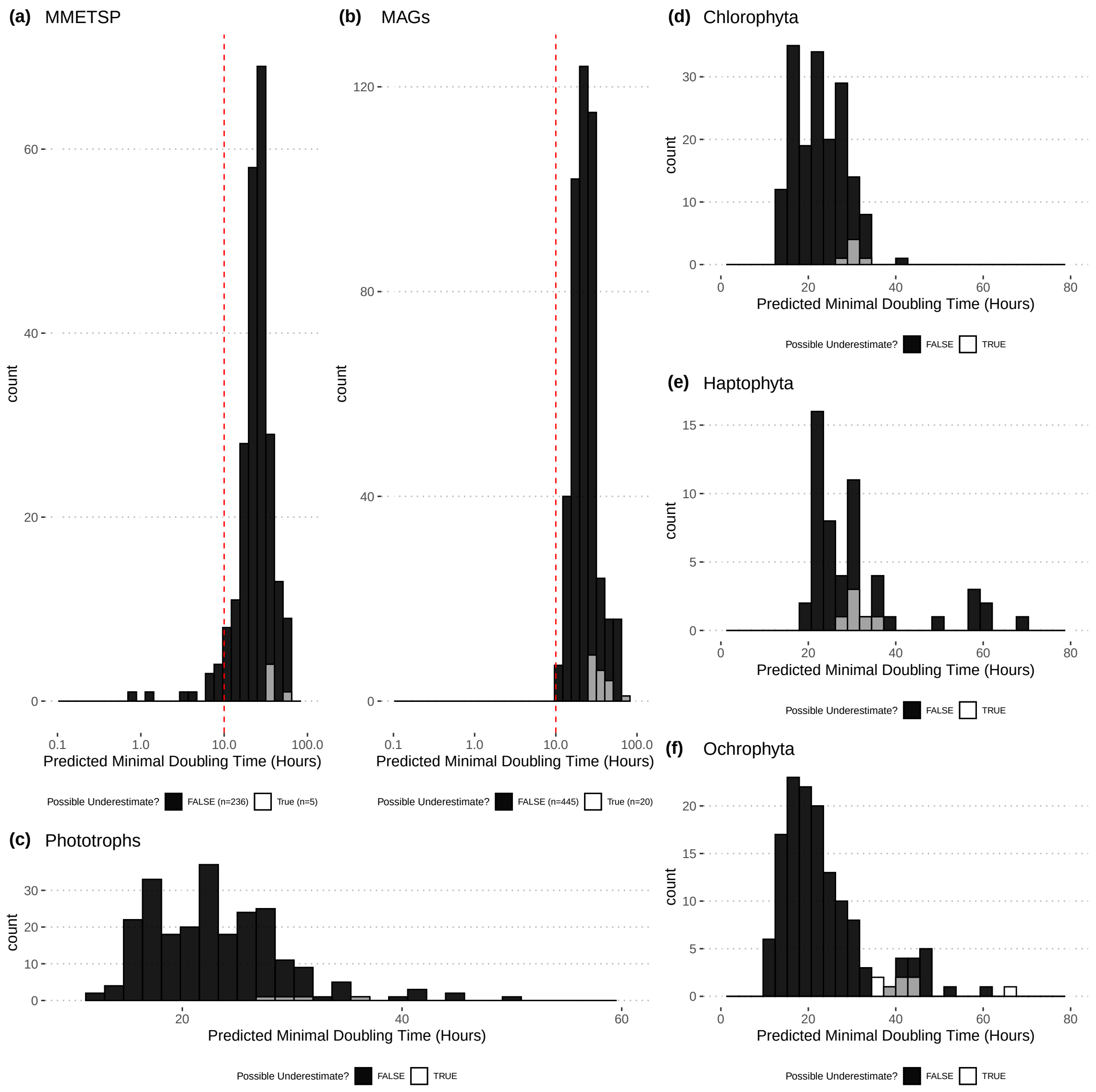


Figure S7: Saturation of the CUB versus maximum growth rate relationship is unlikely to account for the patterns seen in Fig 2. Possible underestimated doubling times determined by codon usage bias cutoff after which the relationship with growth appears to saturate (see Methods).

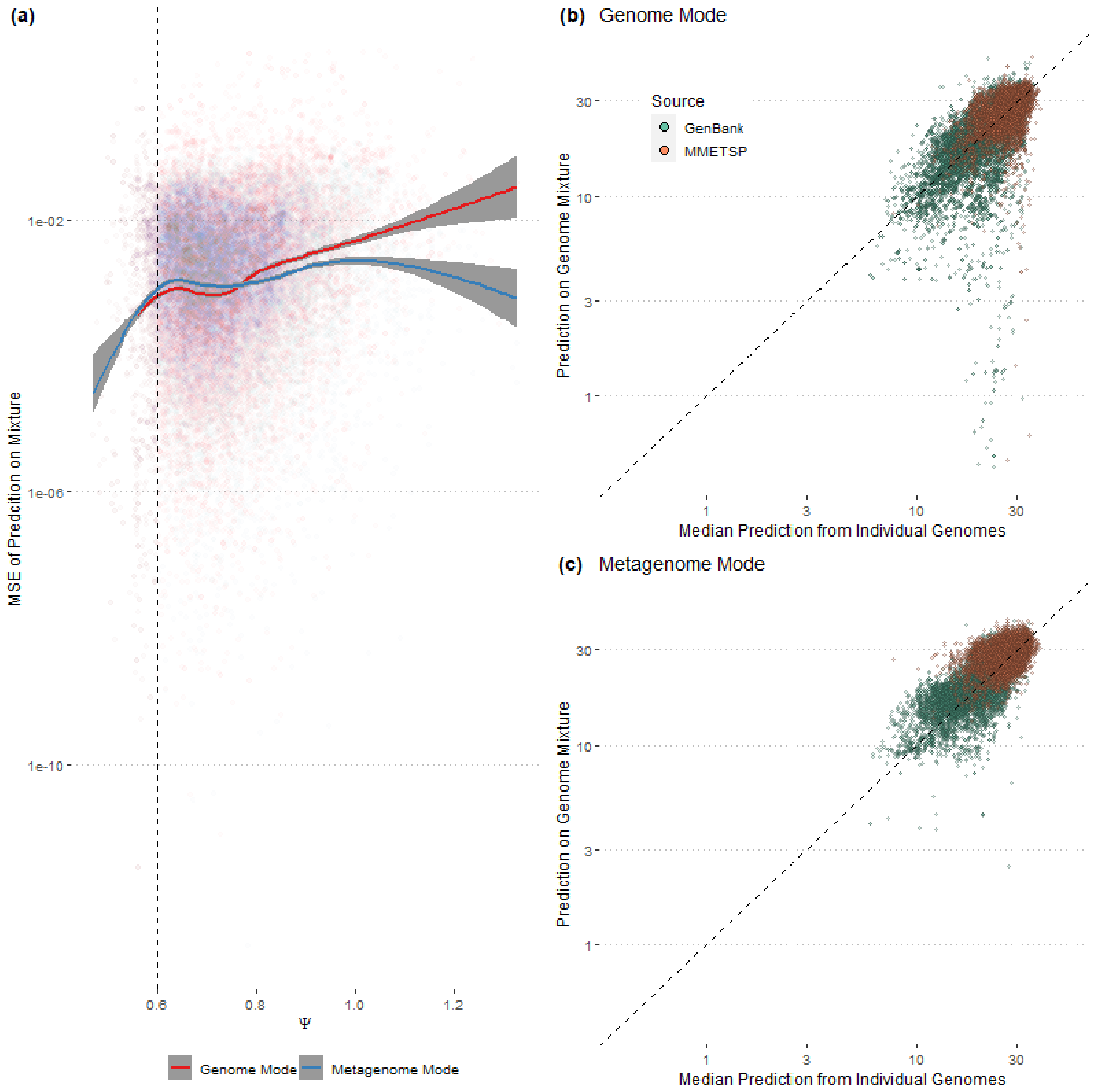


Figure S8: Benchmarking eukaryotic metagenome mode. (a) How well each predictor applied to a gene mixture corresponds to the average of predictions on the individual source genomes (MSE: mean squared error). The value $\Psi$ is a measure of how different the codon usages are of the organisms in a mixture. Note that for high $\Psi$ values (very different codon usages) eukaryotic metagenome mode outperforms eukaryotic genome mode. This can be seen in panel (b) as a tail of organisms falling off-diagonal for genome mode that are close to the diagonal in panel (c).

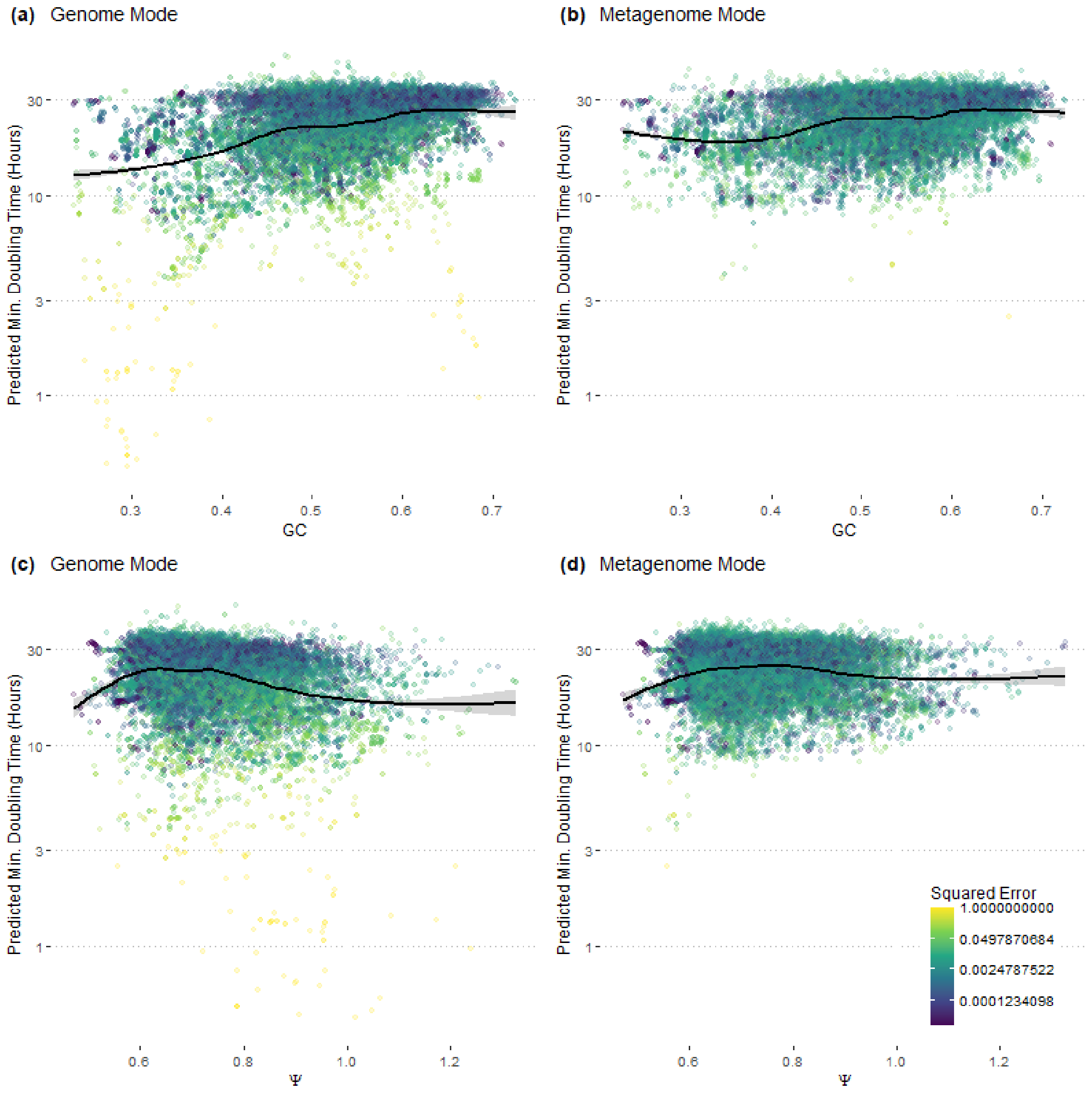


Figure S9: For mixed-species communities eukaryotic genome mode has a slight GC skew and possibly some $\Psi$ skew that is not apparent in metagenome mode. Similar results have been found when benchmarking gRodon predictions on prokaryotic communities.

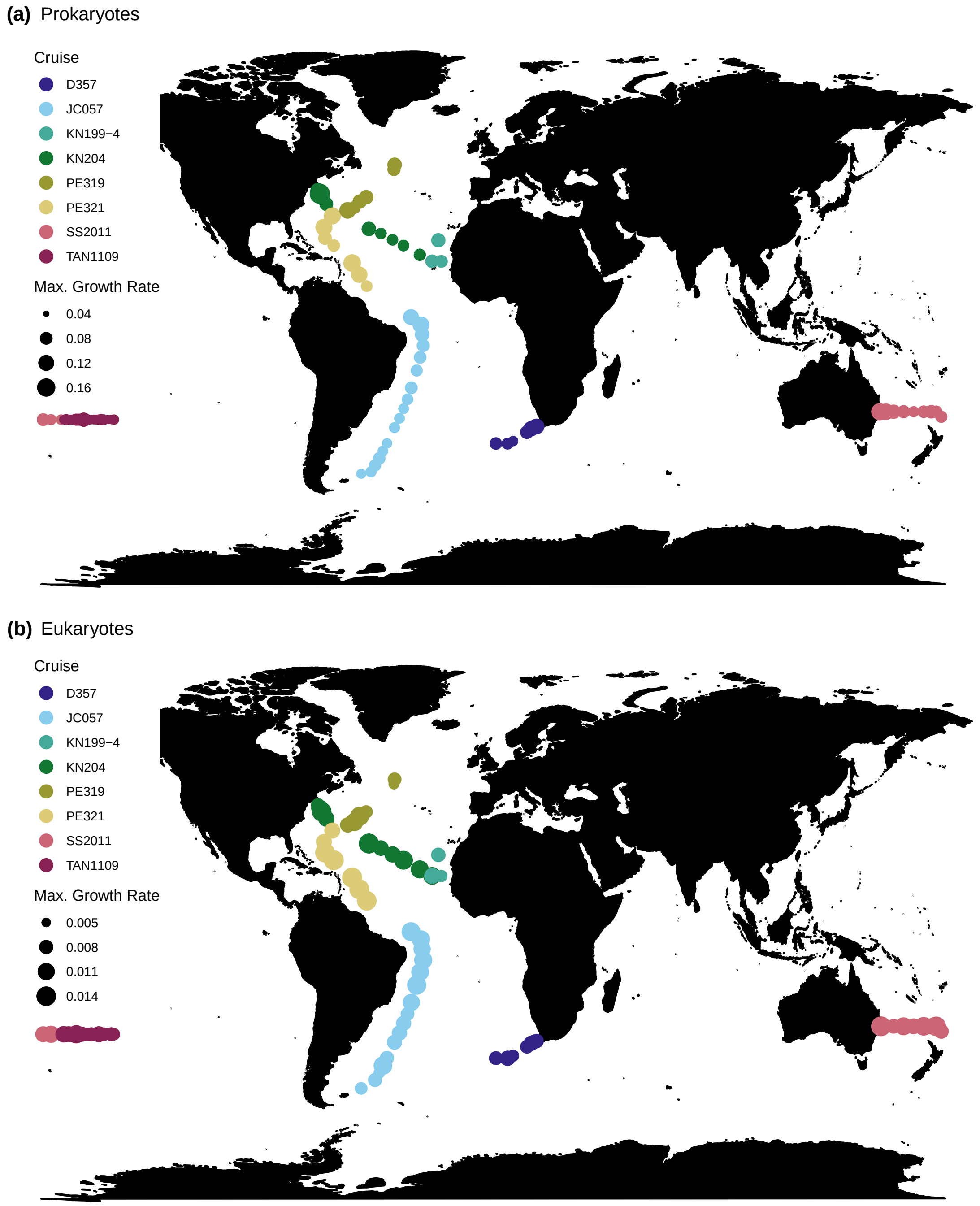


Figure S10: Average community-wide growth rates of BioGEOTRACES surface samples (< 100 meters), broken down by cruise ID.

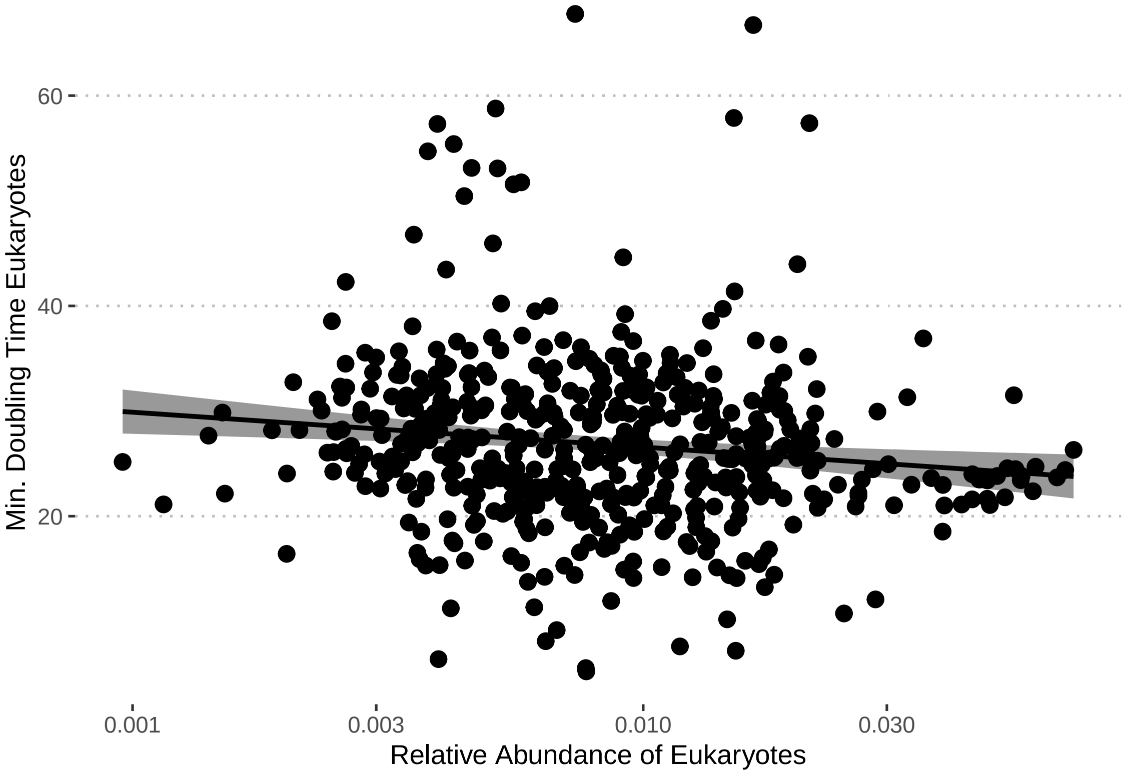


Figure S11: The average minimal doubling time of the eukaryotic community is negatively correlated with the relative abundance of eukaryotic contigs (percent of total reads; Pearson correlation, $p = 0.00414$, $\rho = -0.126$), though the relationship is not particularly strong.

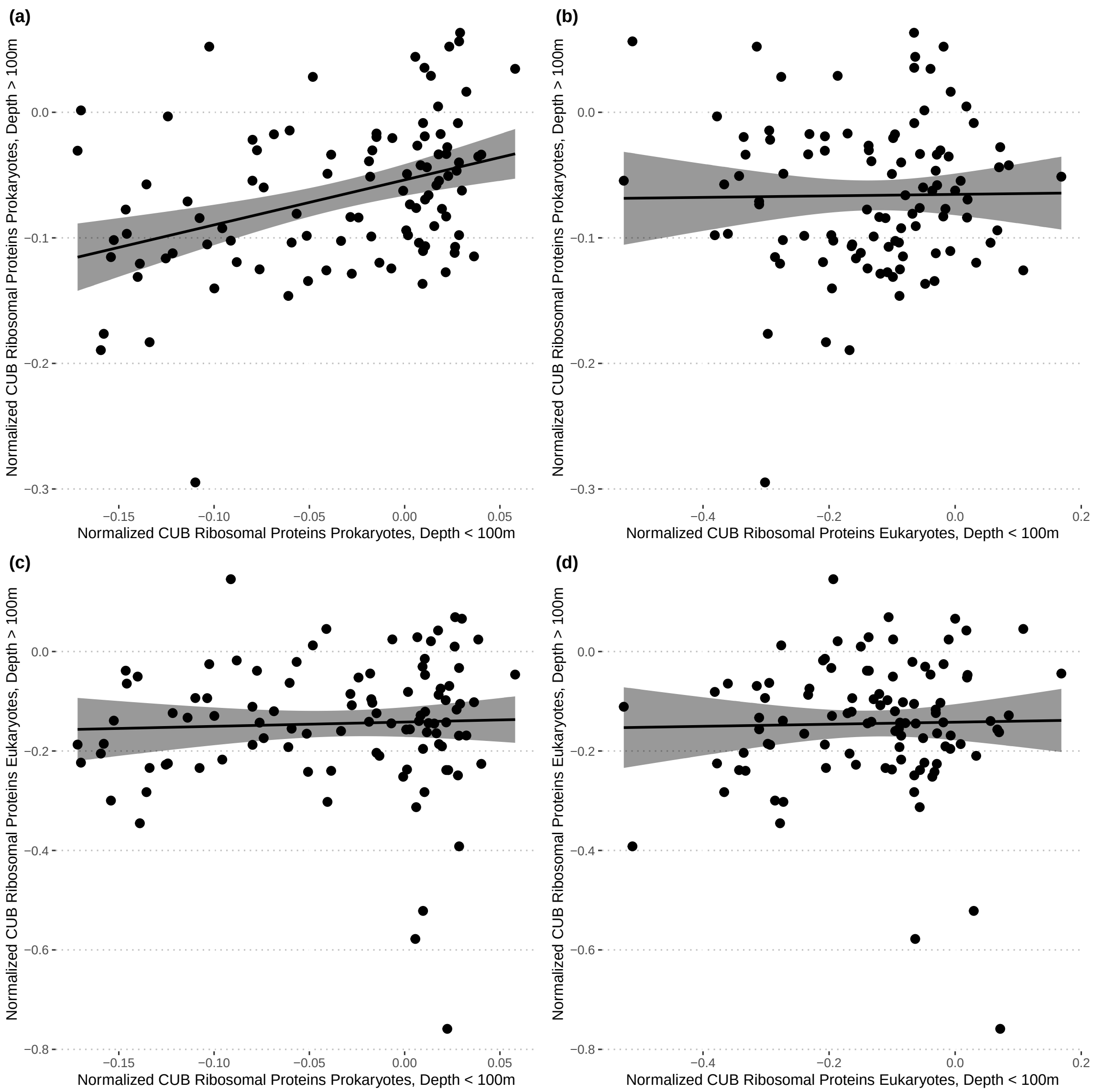


Figure S12: The codon usage biases of eukaryotic and prokaryotic communities are only sometimes correlated across depth. (a) Prokaryotic communities have significant cross-depth correlation (Pearson correlation, $\rho = 0.374$, $p < 1.25 \times 10^{-4}$). (c-e) No other relationships were sigificant.

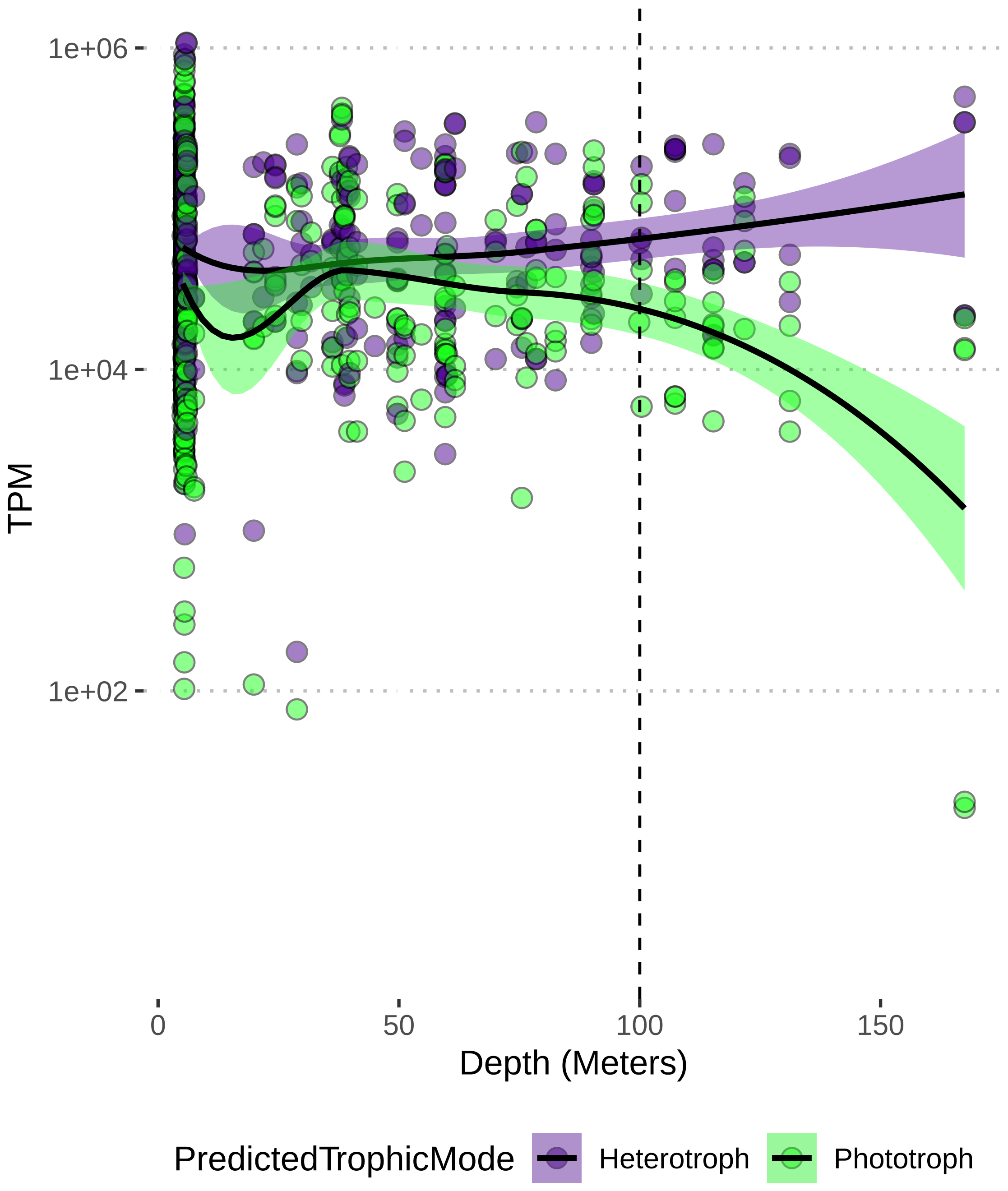


Figure S13: Abundance of the Alexander et al. TOPAZ MAGs across the Tara Oceans metagenomes, where each point represents the sum of heterotrophic or phototrophic MAG relative abundances in a given sample.

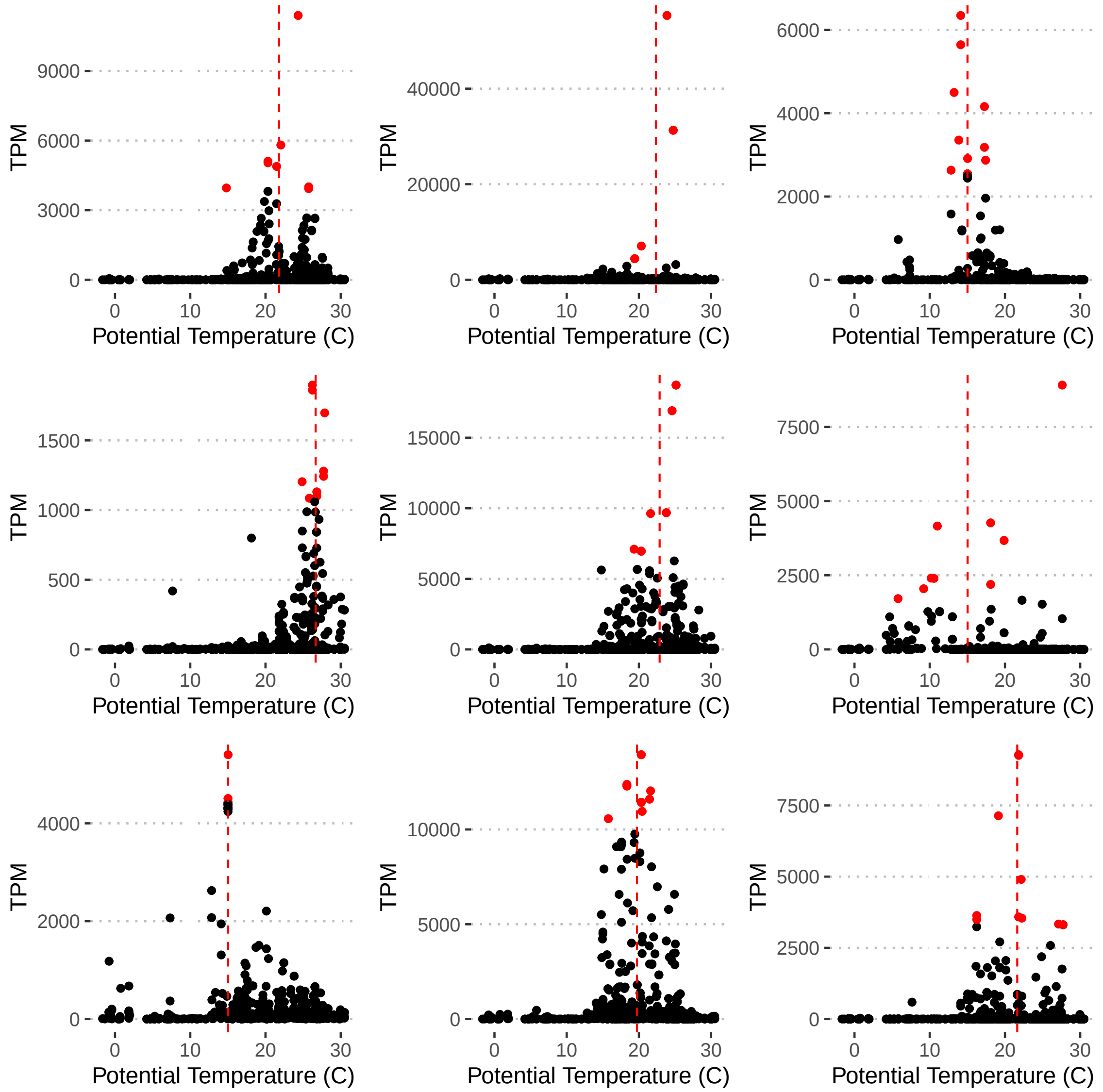


Figure S14: Example relative abundance profiles across Tara Oceans metagenomes for Alexander et al. (TOPAZ) MAGs used to estimate optimal growth temperature. We took the mean temperature (vertical dashed red line) of the samples with the top 1% of relative abundances (red points) among all samples for each MAG. The profiles for 9 MAGs shown were drawn at random from the dataset for visualization of this process.

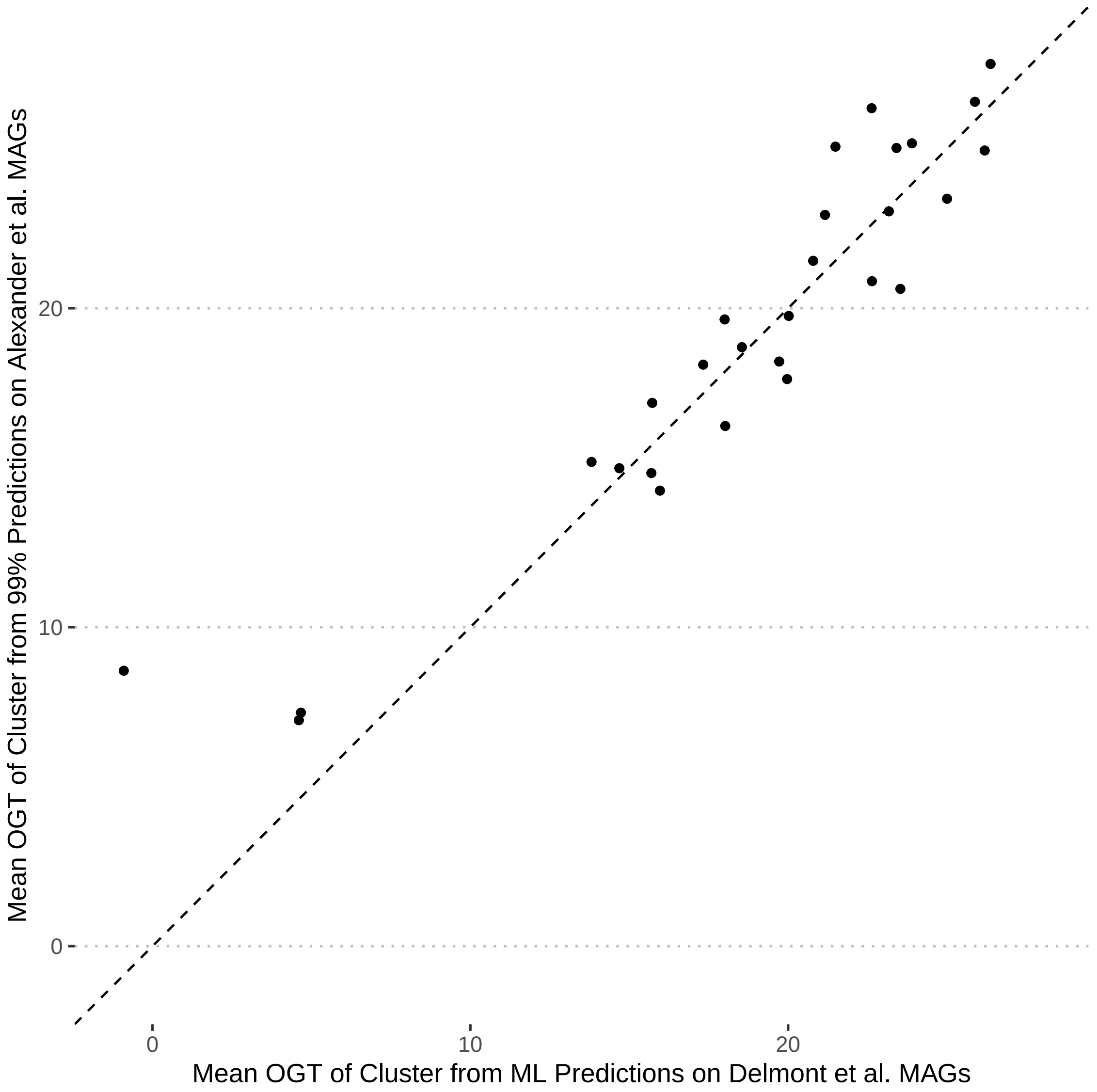


Figure S15: The predictions from the 99th-percentile method used to estimate the optimal growth temperature of the Alexander et al. MAGs agrees well with the machine-learning niche predictions from Delmont et al. (each point represents a cluster of MAG clustered at 99% identity that had both Alexander et al. and Delmont et al. MAGs; mean temperature taken when multiple MAGs from same source were found in the same cluster). The overall correlation between the two estimation methods was high ($\rho = 0.933$, $p = 4.39^{-13}$).

Table S2: Ribosomal proteins retrieved from the Ribosomal Protein Gene Database (`http://ribosome.med.miyazaki-u.ac.jp/|`)

| Species | Ribosomal Protein Genes |
|---|---|
| *Dictyostellium discoideum* | RpsA, Rps2, Rps3, Rps3a, Rps4, Rps5, Rps6, Rps7, Rps8, Rps9, Rps10, Rps11, Rps12, Rps13, Rps14, Rps15, Rps15a, Rps16, Rps17, Rps18, Rps19, Rps20, Rps21, Rps23, Rps24, Rps25, Rps26, Rps27, Rps27a, Rps28, Rps29, Rps30, Rpl3, Rpl4, Rpl5, Rpl6, Rpl7, Rpl7a, Rpl8, Rpl9, Rpl10, Rpl10a, Rpl11, Rpl12, Rpl13, Rpl13a, Rpl14, Rpl15, Rpl17, Rpl18, Rpl18a, Rpl19, Rpl21, Rpl22, Rpl23, Rpl23a, Rpl24, Rpl26, Rpl27, Rpl27a, Rpl28, Rpl29, Rpl30, Rpl31, Rpl32, Rpl34, Rpl35, Rpl35a, Rpl36, Rpl36a, Rpl37, Rpl37a, Rpl38, Rpl39, Rpl40, Rplp0, Rplp1, Rplp2 |
| *Giardia lamblia* | RpsA, Rps2, Rps3, Rps3a, Rps4x, Rps5, Rps6, Rps7, Rps8, Rps9, Rps10, Rps11, Rps12, Rps13, Rps14, Rps15, Rps15a, Rps16, Rps17, Rps18, Rps19, Rps20, Rps21, Rps23, Rps24, Rps25, Rps26, Rps27, Rps27a, Rps28, Rps29, Rps30, Rpl3, Rpl4, Rpl5, Rpl7, Rpl7a, Rpl8, Rpl9, Rpl10, Rpl10a, Rpl11, Rpl12, Rpl13, Rpl13a, Rpl15, Rpl17, Rpl18, Rpl18a, Rpl19, Rpl21, Rpl22, Rpl23, Rpl23a, Rpl24, Rpl26, Rpl27, Rpl27a, Rpl29, Rpl30, Rpl31, Rpl32, Rpl34, Rpl35, Rpl35a, Rpl36, Rpl36a, Rpl37, Rpl37a, Rpl39, Rpl40, Rplp0, Rplp1, Rplp2 |
| *Phaeodactylum tricornutum* | RpsA, Rps2, Rps3, Rps3a, Rps4, Rps5, Rps6, Rps7, Rps8, Rps9, Rps10, Rps11, Rps12, Rps13, Rps14, Rps15, Rps15a, Rps16, Rps17, Rps18, Rps19, Rps20-1, Rps20-2, Rps21, Rps23, Rps24, Rps25, Rps26, Rps27, Rps27a, Rps28, Rps29-1, Rps29-2, Rps30, Rpl3, Rpl4, Rpl5, Rpl6, Rpl7, Rpl7a, Rpl8, Rpl9, Rpl10, Rpl10a, Rpl11, Rpl12, Rpl13, Rpl13a, Rpl14, Rpl15, Rpl17, Rpl18, Rpl18a, Rpl19, Rpl21, Rpl22, Rpl23, Rpl23a, Rpl24, Rpl26, Rpl27, Rpl27a, Rpl28, Rpl29, Rpl30, Rpl31, Rpl32, Rpl34, Rpl35, Rpl35a, Rpl36, Rpl36a-1, Rpl36a-2, Rpl37, Rple37a, Rpl38, Rpl39, Rpl40, Rpl41, Rplp0, Rplp1, Rplp2-1, Rplp2-2 |
| *Plasmodium falciparum* | RpsA, Rps2, Rps3, Rps3A, Rps4, Rps5, Rps6, Rps7, Rps8, Rps9, Rps10, Rps11, Rps12, Rps13, Rps14, Rps15, Rps15a, Rps16, Rps17, Rps18, Rps19, Rps20, Rps21, Rps23, Rps24, Rps25, Rps26, Rps27, Rps27a, Rps28, Rps29, Rps30, Rpl3, Rpl4, Rpl5, Rpl6, Rpl7, Rpl7a, Rpl8, Rpl9, Rpl10, Rpl10a, Rpl11, Rpl12, Rpl13, Rpl13a, Rpl14, Rpl15, Rpl17, Rpl18, Rpl18a, Rpl19, Rpl21, Rpl22, Rpl23, Rpl23a, Rpl24, Rpl26, Rpl27, Rpl27a, Rpl28, Rpl29, Rpl30, Rpl31, Rpl32, Rpl34, Rpl35, Rpl35a, Rpl36, Rpl36a, Rpl37, Rpl37a, Rpl38, Rpl39, Rpl40, Rpl41, Rplp0, Rplp1, Rplp2 |
| *Thalassiosira pseudonana* | RpsA, Rps2, Rps3, Rps3a, Rps4, Rps5, Rps6, Rps7, Rps8, Rps9, Rps10, Rps11, Rps12, Rps13, Rps14, Rps15, Rps15a, Rps16, Rps17, Rps18, Rps19, Rps20, Rps21, Rps23, Rps24, Rps25, Rps26, Rps27, Rps27a, Rps28, Rps29, Rps30, Rpl3, Rpl4, Rpl5, Rpl6, Rpl7, Rpl7a, Rpl8, Rpl9, Rpl10, Rpl10a, Rpl11, Rpl12, Rpl13, Rpl13a, Rpl14, Rpl15, Rpl17, Rpl18, Rpl18a, Rpl19, Rpl21, Rpl22, Rpl23-1, Rpl23-2, Rpl23a, Rpl24, Rpl26, Rpl27, Rpl27a, Rpl28, Rpl29, Rpl30, Rpl32, Rpl34, Rpl35, Rpl35a, Rpl36, Rpl36a, Rpl37, Rpl37a, Rpl38, Rpl39, Rpl40, Rpl41, Rplp0, Rplp1, Rplp2 |
| *Toxoplasma gondii* | RpsA, Rps2, Rps3, Rps3a, Rps4, Rps5, Rps6, Rps7, Rps8, Rps9, Rps10, Rps11, Rps12, Rps13, Rps14, Rps15, Rps15A, Rps16, Rps17, Rps18, Rps19, Rps20, Rps21, Rps23, Rps24, Rps25, Rps26, Rps27, Rps27a, Rps28, Rps29, Rps30, Rpl3, Rpl4, Rpl5, Rpl6, Rpl7, Rpl7a, Rpl8, Rpl9, Rpl10, Rpl10a, Rpl11, Rpl12, Rpl13, Rpl13a, Rpl14, Rpl15, Rpl17, Rpl18, Rpl18a, Rpl19, Rpl21, Rpl22, Rpl23, Rpl23a, Rpl24, Rpl26, Rpl27, Rpl27a, Rpl28, Rpl29, Rpl30, Rpl31, Rpl32, Rpl34, Rpl35, Rpl35a, Rpl36, Rpl36a, Rpl37, Rpl37a, Rpl38, Rpl39, Rpl40, Rpl41, Rplp0, Rplp1, Rplp2 |
